# Tracking the zombie ant graveyards’ dynamics at Mulu National Park, Borneo, Malaysia

**DOI:** 10.64898/2025.12.18.695298

**Authors:** Daniel Chandler, Jack Williams, Chayli McCann, Kimberley Hui Li Tan, Zoe Irvine, Anushkaa Lakmani Vasu, Eliza Planincic, Sze Huei Yek

## Abstract

*Ophiocordyceps unilateralis sensu lato* is a parasitoid fungus that predominantly infects ants from the Camponotini tribe, inducing behavioural changes to enhance its survival and reproduction. The most notable change occurs during the final infection phase, where the infected ant leaves its nest to perform the ‘death grip’, hanging upside down and biting onto a substrate to anchor itself. In the tropics, these infected ants congregate at specific locations known as zombie graveyards, which have optimal conditions for spore dispersal. This study examines the spatial dynamics of known graveyards at a national park in Borneo, Malaysia, by assessing existing sites and identifying new ones. We located old graveyards at Night Walk and Paku Trail and discovered a new one at Botany Loop. The density of ant cadavers per quadrat increased slightly, from 1-3 ants/m^2^ to 2-4 ants/m^2^. While the graveyard size at Night Walk remained similar, Paku Trail experienced a nearly two-fold reduction, decreasing from 197 m^2^ to 97 m^2^. At Night Walk, three ant hosts were identified, with *Colobopsis leonardi* being the dominant ant host across all graveyards. Without genetic tests, we could not confirm whether the same *O. unilateralis* infects all ants in the graveyards. Our findings suggest a general site fidelity of zombie graveyards, with hotspot densities varying spatially, potentially influenced by microclimatic changes. Long-term monitoring of graveyard dynamics is essential for understanding natural infections and the effects of environmental conditions on parasitoid-host ecology.

## Introduction

*Ophiocordyceps unilateralis sensu lato* fungi is a pantropical species-complex that parasitises ants from the Camponotini tribe (Lin et al. 2020; Araújo et al. 2018; 2020). The fungi induce physiological and behavioural changes in their hosts, supporting parasite development, reproduction, and transmission (Andriolli et al. 2019; Lefevre et al. 2009). The fungal parasites infect forager ants with spores that attach to and penetrate the ant’s exoskeleton (Andriolli et al. 2019). The suite of behavioural changes in ant hosts induced by the fungal parasite is termed the ‘extended phenotype’ of the fungi (Andersen et al. 2009). The behavioural changes are due to the activation of particular genes within the ant host and parasite (de Bekker and Das 2022). The distinctive behavioural anomalies exhibited by infected ants occur in two phases. Within their arboreal nests, infected ants abandon routine behaviours and begin to walk randomly while convulsing periodically (Beckerson et al. 2023). This causes the ants to fall from their nest (Hughes et al. 2011). Once the infected ant is on the forest floor, a set of behaviours known as ‘summit disease’ manifests (Andriolli et al. 2019). Random walking is replaced by deliberate movement, which responds to specific environmental cues. The infected ant moves towards locations favourable for fungal growth (Anderson et al. 2009; de Bekker 2019). Once it reaches the location, the ant bites onto a substrate (e.g. small twigs or on the leaf vein) with an unbreakable ‘death grip’ and remains in this position until deceased (Mangold et al. 2019). The fungus grows within the ant’s body for several days and uses the host’s nutrients to support its development. The fruiting body eventually emerges from the base of the ant’s head. The fruiting body releases spores into the surrounding environment onto foraging ants, completing the fungal life cycle (Mangold et al. 2019; Mongkolsamrit et al. 2012).

The infected ants often congregate in the same area, called the ‘zombie graveyards’ (Pontoppidan et al. 2009). The formation of graveyards within rainforests is limited to a small area and patchy in distribution, with occasional clusters of high ant density just metres away from areas with no infected ants (Anderson et al. 2012; Pontoppidan et al. 2009; Lavery et al. 2021). For example, Lavery et al. (2021) found 13 ants per square meter, while Pontoppidan et al. (2009) detected 26 ants per square meter adjacent to quadrats with no infected ants. Over time, the approximate location of the graveyard remains stable (i.e. site fidelity). However, high-density areas in the Thai rainforest became low-density (Anderson et al. 2012; Pontoppidan et al. 2009), suggesting that zombie ant graveyards could be spatially dynamic, with areas of highest infection shifting frequently. These high-density areas (i.e., graveyards) are typically found in the rainforest understory, within a metre above the ground, where moderate canopy shading occurs (Hughes et al. 2011). Graveyards have a unique microclimate that supports the successful growth of fungi. So far, humidity and canopy openness (i.e., light exposure) have been identified as essential factors for *O. unilateralis sensu lato* to produce fruiting bodies (Andriolli et al. 2019; Neto et al. 2019; Will et al. 2023). For example, infected ant cadavers relocated to environments with alternative microclimates failed to sporulate (Andersen et al. 2009; Evans et al. 2011).

*O. unilateralis sensu lato* predominantly infects *Colobopsis leonardi* (formerly *Camponotus leonardi*) (Ward and Boudinot 2021), but other Camponotini ants, such as *Polyrhachis* sp., were sometimes found at the same graveyard with low prevalence (Pontoppidan et al. 2009; Lin et al. 2020). The detection of multiple ant hosts could suggest that there are several cryptic species within *O. unilateralis*. This finding is consistent with those of Evans et al. (2011), who discovered diversity within *O. unilateralis* across four different *Camponotus* species. The currently accepted notion is that *O. unilateralis* is a species complex with noticeable local diversities of associated ant hosts. However, in an extensive survey conducted in 2019 at four graveyards in Mulu National Park, Sarawak, Borneo, Malaysia, only *C. leonardi* ant hosts were detected (Lavery et al. 2021).

Studies suggest zombie graveyards display general site fidelity with shifting hotspots (i.e., areas with high densities of ant cadavers). However, zombie graveyards have not been monitored over the long term (i.e., four years). Hence, this study was set up to achieve this objective. We attempted to locate previous zombie graveyards found in 2019 and to monitor the dynamics inside the graveyards. This assesses whether the densities have changed. We also expanded our survey to cover wider areas and new trails within the Mulu National Park to evaluate whether *O. unilateralis sensu lato* infects other ant hosts.

## Methodology

### Site description

This study was conducted in a tropical dipterocarp forest within Gunung Mulu National Park in Sarawak, Borneo, Malaysia (approximately 528 km^2^), at an altitude of 1845 above sea level. The study was conducted from September 28th to 30th, 2023, at the beginning of the wet season. The time and conditions of the survey were comparable to those of the 2019 zombie ants graveyard survey by Lavery et al. (2021). Their study was conducted from September 26th to 27th. Temperatures in September average a maximum of 28 °C and a minimum of 18 °C. The average precipitation for September was 205 mm (https://www.meteoblue.com/en/weather/historyclimate/climatemodelled/gunung-mulu-national-park_malaysia_6942283).

Zombie ant graveyards were discovered at two trails (i.e., Paku Trail and Night Walk) in 2019 (Fig. 1). To investigate whether the graveyards had shifted, we systematically surveyed the previous sites and also searched for new graveyards. We rediscover graveyards at Paku Trail and Night Walk over the three days. Additionally, we discovered a new graveyard at Botany Loop (Fig. 1).

**Figure 1.**
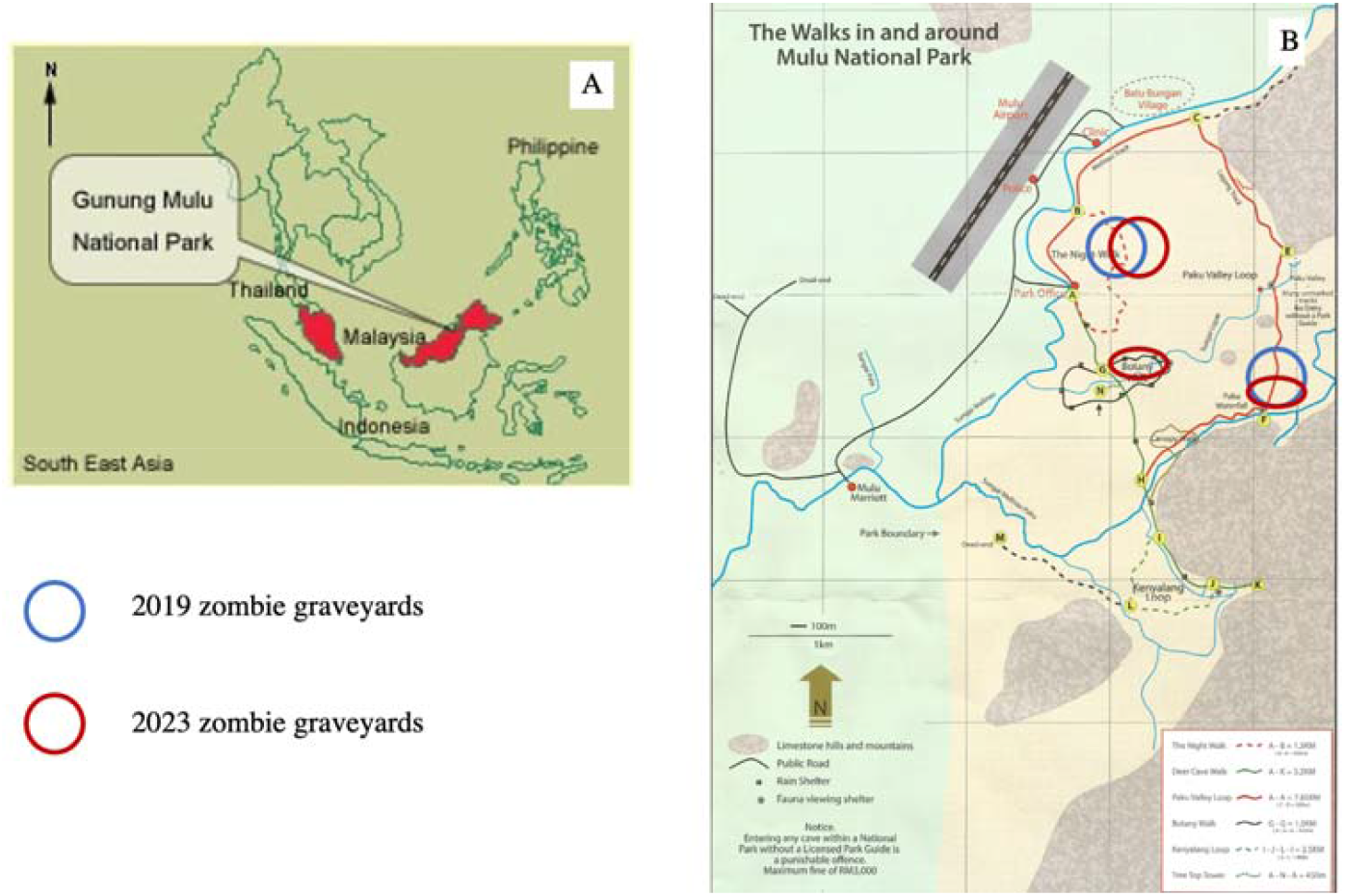
(A) Map of Mulu National Park in Borneo, Malaysia. (B) The 2019 ant graveyards were located at Paku Trail and Night Walk (dark blue circles), whereas 2023 ant graveyards were located at Paku Trail, Night Walk, and a new graveyard was discovered at Botany Loop (dark red circles). Maps were downloaded from the official Mulu National Park webpage (https://mulupark.com/).

### Data Collection

The zombie ant cadaver survey largely follows the search strategy of Lavery et al. (2021): the first infected ant encountered at the 2019 graveyard was selected as the starting point of each transect, ensuring that subsequent sampling activities were conducted within the confines of the ant graveyard. Instead of sampling in four directions (North, South, East, West) of the starting 1 m^2^ quadrat, we conducted searches along a transect line starting from the first ant cadaver quadrat in only two directions, up and down the trail where the ant cadavers were found. Five quadrats (1 m^2^ x 1 m^2^ per quadrat) were set up at 10 m intervals along the transect line. Each observer was responsible for five quadrats. This modification to the search strategy aimed to cover a broader area to better identify the extent of the graveyards.

Zombie ant cadaver searching entailed visually inspecting the surface of all plant leaves and small twigs up to a height of about two metres within each quadrat. Leaves were gently turned over and assessed for the presence of an ant cadaver underside the leaf. Zombie ant cadavers can sometimes occur at heights above two metres, but most ant cadavers were confined within a metre above ground at this location (Lavery et al. 2021). Accurate counting above two metres necessitated equipment (i.e., ladder or tree climbing gear), which was not available for this study. Once an ant cadaver was detected, the location was demarcated with coloured tape, facilitating the subsequent quantification of the total number of ant cadavers within each quadrat. At least one intact ant cadaver was collected from each quadrat for host ant identification confirmation. Any infected ants that appeared morphologically different from *Colobopsis leonardi* were also collected for further examination.

The search for ant cadavers continued along the transect line at each trail until the absence of ant cadavers in all five quadrats within the 10-metre transect interval in both directions, marking the endpoints of the graveyard. A measuring tape was then used to determine the total distance between the two endpoints, enabling an approximation of the graveyard’s area. The number of infected ants identified within each quadrat was counted, yielding cumulative totals for each quadrat (Supplementary Table 1). Ant cadavers with fungal growth underwent microscopic examination to enable taxonomic identification to the lowest taxonomic level. As the research permit only covered observational data, photographic documentation of the ant cadavers was obtained, and the trails from which they were collected were recorded.

**Table 1:** Summary of graveyard characteristics at three trails in Gunung Mulu National Park, Borneo, Malaysia. Graveyards were found on Night Walk and Paku Trail in 2019 and rediscovered in 2023. A new graveyard was found on Botany Loop in 2023.

|  | Night Walk<br>(2019) | Night Walk<br>(2023) | Paku Trail<br>(2019) | Paku Trail<br>(2023) | Botany<br>Loop<br>(2019) | Botany<br>Loop<br>(2023) |
| --- | --- | --- | --- | --- | --- | --- |
| Number of<br>Ants | 292 | 114 | 255 | 21 | - | 42 |
| Graveyard<br>area (m<sup>2</sup>) | 299 | 272 | 191 | 97 | - | 102 |
| Mean ant<br>cadavers<br>per quadrat<br>(ants/m<sup>2</sup>) | 1 | 4 | 1 | 2 | - | 3 |
| Number of<br>infected<br>host ants | 1 | 3 | 1 | 1 | - | 1 |

### Data Analysis

Data collation was performed using Microsoft Excel, and subsequent analysis was conducted in RStudio. A descriptive comparison was conducted between the graveyards in 2019 (Lavery et al. 2021) and 2023. Since the sampling strategy differed from that of Lavery et al. (2021), statistical comparisons between years were not possible. Instead, we compare the surface area of the graveyards and the density of ant cadavers found per quadrat. A Pearson correlation was run to determine the relationship between graveyard size and the number of ant cadavers found.

## Results

### Graveyard dynamics over five years

Lavery et al. (2021) identified four graveyard sites, three at Night Walk and one at Night Paku Trail. The current study categorises all dead ant cadavers on the same trail belonging to the same graveyards. Hence, for comparison purposes, we combined the three graveyard sites found on the Paku Trail in 2019 as a single graveyard in this study (Table 1). The current sampling strategy aims to locate the extent of the graveyard size and its overall spatial dynamics; hence, the sampling strategy differs slightly. For 2019, two successive empty quadrats in all directions demarcate the graveyard’s edge. In contrast, the current study demarcates five empty quadrats 10 metres away from the last ant cadavers as the graveyard’s edge.

The graveyard size on Night Walk was similar in 2019 and 2023, at 299 m^2^ and 272 m^2^, respectively (Table 1). However, the Paku Trail graveyards decreased substantially from 191 m^2^ in 2019 to 97 m^2^ in 2023. A new graveyard was found on Botany Loop. No graveyard was detected at Botany Loop in the 2019 surveys. The mean density of ant cadavers found per quadrat increased from 1 to 3 ants/m^2^ in 2019 to 2 to 4 ants/m^2^. From the Pearson correlation, we found a moderate, positive correlation between graveyard size and the number of ant cadavers, which is not statistically significant (*r*(4) = 0.800, *p* = 0.055).

### Multiple ant hosts at the Night Walk graveyard

Only one ant host, *Colobopsis leonardi*, was found in the 2019 study (Lavery et al. 2021). In the 2023 survey, we found three different ant hosts at Night Walk (Table 1; Fig. 2). *Colobopsis leonardi* was the dominant ant host, with the other two hosts (both identified as *Camponotus* spp.) comprising 7.89% (N=9) of the ant cadavers found in the Night Walk graveyard. For graveyards on Paku Trail and Botany Loop, only *C. leonardi* ant hosts were found.

**Figure 2.**
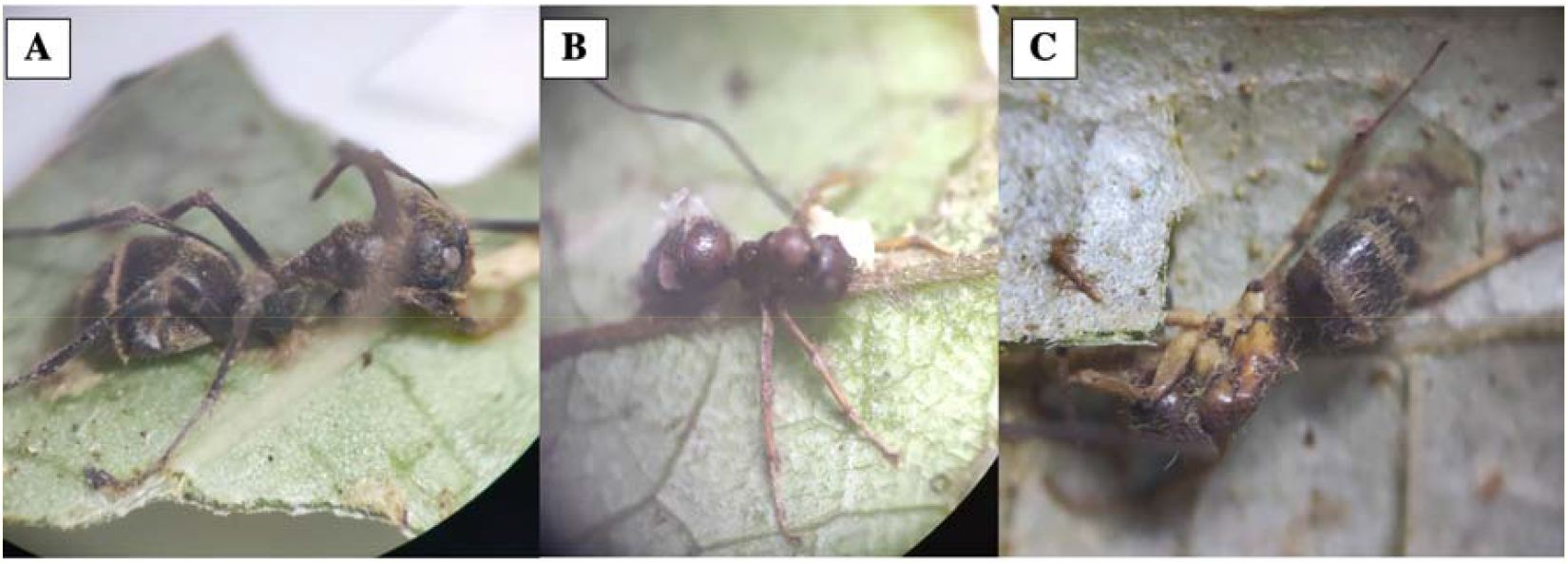
Three ant hosts found on Night Walk graveyard. A is *Colobopsis leonardi*, B is *Camponotus* sp. B and C is *Camponotus* sp. C.

*Colobopsis leonardi* is characterised by golden hairs on its head, mesosoma, and gaster. This host ant is characterised by uniform dark brown or black legs (Fig. 2A). *Camponotus* sp. B’s head was dark brown with no visible hairs and had a shorter pronotum than *C. leonardi*. The gaster displayed medium to long, uniformly distributed hair with light brown leg colour (Fig. 2B), and there were eight *C*. sp. B’s cadavers at Night Walk. *Camponotus* sp. C had a dark brown head, a light brown mesosoma, striped hair formation on the gaster, and legs featuring a mix of yellow and black colour (Fig. 2C), and there was only one *C*. sp. C’s cadaver at Night Walk (Supplementary Table 1).

## Discussion

Field examination of graveyards indicates a general site fidelity (Pontoppidan et al. 2009; Anderson et al. 2012; Will et al. 2023). However, these insights were gained from tracking graveyards for one to two years. Our study represents the longest (i.e., four years) site fidelity for graveyard locations. In 2019, graveyards were discovered on two trails (Night Walk and Paku Trail) (Lavery et al. 2021). In 2023, these graveyards were re-discovered, and a new graveyard was located on a new trail (Botany Loop) at Mulu National Park, Borneo, Malaysia. We cannot exclude the Botany Loop graveyard as a pre-existing graveyard missed in the 2019 survey. Abiotic factors, such as canopy openness and humidity, are important drivers in graveyard locations (Andriolli et al. 2019; Neto et al. 2019; Will et al. 2023). The site fidelities displayed by these continuously infected ant cadavers suggest that changes in graveyard locations are constrained by the availability of suitable abiotic conditions, resulting in the recurrent use of the locations over extended periods.

The formation of graveyards is often limited to a small area and patchy in distribution, with occasional clusters of high density just metres away from areas with no infected ants (Anderson et al. 2012; Pontoppidan et al. 2009; Lavery et al. 2021). Over time, we would expect microclimatic changes at the general graveyard’s location, necessitating spatial shifts in areas with the highest infection rates (Pontoppidan et al. 2009). Our study monitors the graveyard size and density of ants/m^2^ over four years to assess how graveyard dynamics vary. In general, the larger the graveyard size, the more ant cadavers can be found. However, this is not statistically significant. One graveyard (Night Walk) size remains similar, but the density of infected ants increased from 1 ant/m^2^ to 4 ants/m^2^. In contrast, the Paku Trail graveyard size underwent a twofold reduction, whereas the ants’ density increased from 1 ant/m^2^ to 2 ants/m^2^. In the current study, the methodology was modified to cover a wider area; hence, direct statistical comparison is not possible. However, from the reduced graveyard size and increased ant cadaver density, we can infer that there is perhaps a scarcity of suitable microclimatic patches for optimal fungal development in 2023 compared to 2019.

*Ophiocordyceps unilateralis sensu lato* predominantly infects *Colobopsis leonardi* (formerly *Camponotus leonardi*) (Ward and Boudinot 2021), but other Camponotini ants can be found at the same graveyard with low prevalence (Pontoppidan et al. 2009; Lin et al. 2020). The 2019 survey only locates *C. leonardi* in graveyards (Lavery et al. 2021). However, our 2023 survey located two other Camponotini ant hosts sharing the same graveyard with the dominant *C. leonardi*. Detecting multiple ant hosts in the same graveyard could indicate that several cryptic *O. unilateralis* species share the same microclimatic conditions for optimal fungal growth (Evans et al. 2011). However, without further genetic testing, we could not determine whether the *O. unilateralis* infecting the other Camponotini ant hosts belonged to the identical genotypes as the *O. unilateralis* infecting *C. leonardi*.

## Conclusion

This is the first long-term field tracking of zombie graveyards. We found general site fidelity that varies spatially, most probably tracking the microclimatic changes in the graveyards. We also observed multiple ant hosts in one of the graveyard sites, suggesting the presence of cryptic species among *O. unilateralis* that share the same graveyard. We think these graveyards should be reassessed every few years, as longer-term data are valuable for studying natural infections and how environmental conditions affect parasitoid-host ecology.

## Acknowledgements

This study was conducted at Monash University as part of the unit ‘Tropical Terrestrial Biology’, and we wish to recognise the university for providing us with the means to undertake this research. We also thank Gunung Mulu National Park management for permitting us to conduct this study within the park (Permit Number: SFC.810-4/6/1 (2023) - 149).

## Notes

### Competing Interest Statement

The authors have declared no competing interest.

